# One-sided design of protein-protein interaction motifs using deep learning

**DOI:** 10.1101/2022.03.30.486144

**Authors:** Raulia Syrlybaeva, Eva-Maria Strauch

## Abstract

Protein-protein interactions are part of most processes in life and thereby the ability to generate new ones to either control, detect or inhibit them has universal applications. However, to develop a new binding protein to bind to a specific site at atomic detail without any additional input is a challenging problem. After DeepMind entered the protein folding field, we have seen rapid advances in protein structure predictions thanks to the implementation of machine learning algorithms. Neural networks are part of machine learning and they can learn the regularities from their input data. Here, we took advantage of their capabilities by training multiple neural networks on co-crystal structures of natural protein complexes. Inspired by image caption algorithms, we developed an extensive set of NN-based models, referred to as iNNterfaceDesign. It predicts the positioning and the secondary structure for the new binding motifs and then designs the backbone atoms followed by amino acid sequence design. Our methods are capable of recapitulating native interactions, including antibody-antigen interactions, while they also capable to produce more diverse solutions to binding at the same sites. As it was trained on natural complexes, it learned their features and can therefore also highlight preferential binding sites, as found in natural protein-protein interactions. Our method is generally applicable, and we believe that this is the first deep learning model for one-sided design of protein-protein interactions.

**Abstract figure:** 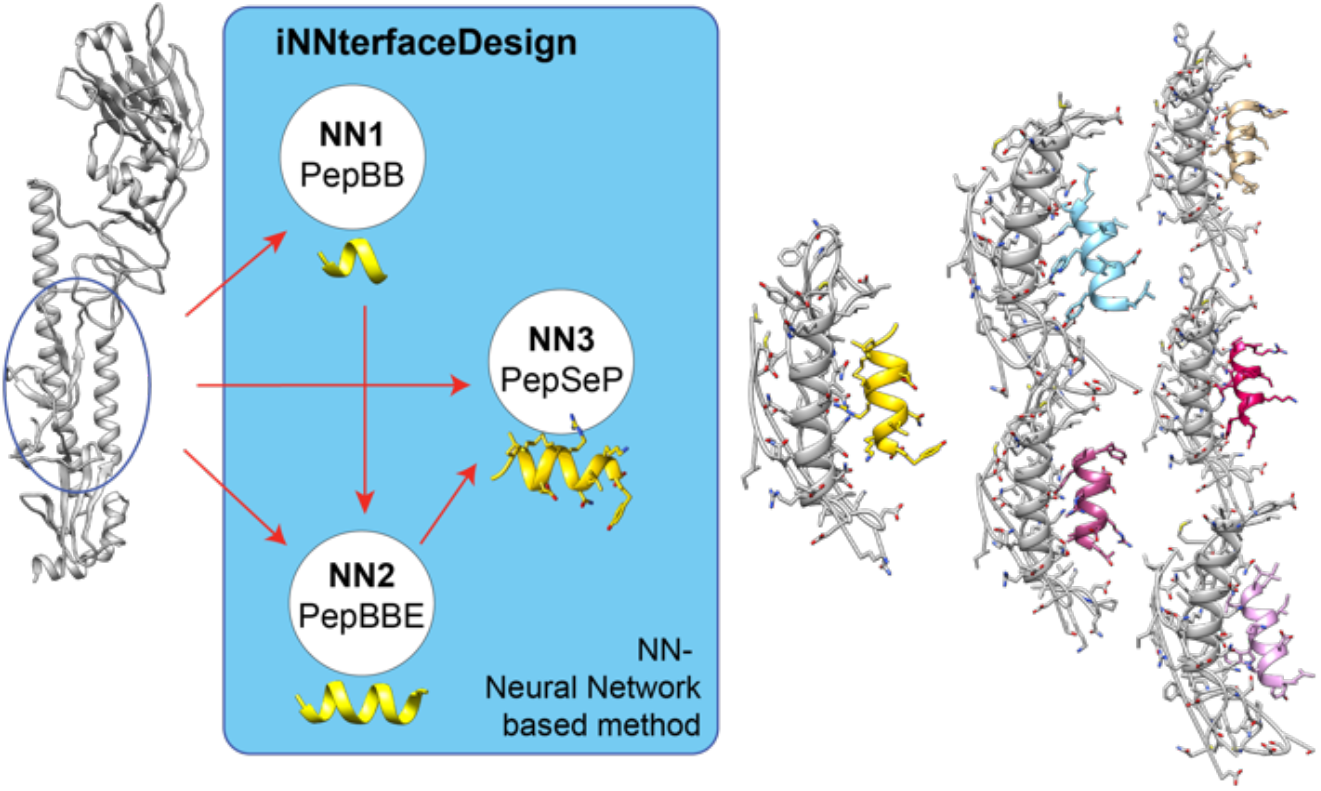

## Introduction

The challenges for the *de novo* design of protein-protein interfaces is to align or design from scratch a new protein backbone to not only fit seamlessly onto the targeted site like a jigsaw puzzle piece, but furthermore, the backbone has to present perfectly a few select contacts that can provide the majority of the binding energy. The problem is that a typical protein-protein interface consists of uneven contacts with only a few “hotspot” interactions which provide the “glue” between the two proteins. Previous approaches to design protein interactions from scratch have used a dock-design approach^1,2^ in which the given protein or an all alanine helix is repetitively docked against a previously identified interface of the target protein and then redesigned using the RosettaDesign^3^ hoping that a compatible docking conformation would allow the design of hotspots. Another approach has been to focus on hotspots directly by starting with disembodied hotspots. However, hotspots were identified from co-crystal structures and then re-docked as an individual single amino acid residue to diversify its orientation, thereby allowing different backbones (and thus proteins) to present them^4–6^. This search had several shortcomings; it was limited to only certain backbone conformations; sampling of hotspot residues positioning was poorly sampled and many redocked residues had none-productive backbone orientations which increased the challenge to identify protein backbones that could harbor these residues. In a more recent advancement^7^, an exhaustive, hierarchical docking search of single, disembodied residues was performed using a simplified forcefield, and scaffolds were then placed onto these to host these contacts, regardless of their binding energies. In a second iterations, backbones fragments presenting some of the pre-docked residues were extracted and re-grafted into more scaffold proteins. The remainder of the interface was then redesigned as part of this focused search. While this approach resulted in the successful design of new binding proteins against 12 different targets, it required both massive computing resources and experimental screening of 715,000 designs^7^. This highlights that the problem is tremendously challenging, but solvable.

Here, we developed a fundamentally new algorithm, referred to as iNNterfaceDesign, for the *de novo* design of protein-protein interfaces. Our method generates *de novo* interface fragments (interaction motifs) based on properties of the target protein alone using several neural networks. The algorithm consists of several neural network (NN) which interface with the Rosetta modeling software^8^ for refinement of the generated protein fragments. Thus, we are integrating deep learning with biophysical modeling. Our algorithms are motivated by the computer vision field which is one of the most developed areas within machine learning. It is inspired by NN-based learning models for the generation of captions for images using visual attention^9^. There are a few examples of successful implementation of pattern recognition techniques in biochemistry, including development of models for the prediction of protein structures^10–13^ based on primary amino acid sequences, such as AlphaFold^14,15^ or RoseTTAFold^16^. Further, new NN-based methods are being developed to design protein folds, either on sequence-based models^17^, ‘hallucination’^18^ or using learned potentials^19^. However, thus far, to our knowledge, there is no deep learningbased design method for the development of PPIs.

Our here discussed iNNterfaceDesign pipeline is able to reproduce interface motifs of user-defined length deviating from interface fragments of native PPIs with root mean square deviation (RMSD) of 3.0-5.5 Å in average for the central 6 residues. The motifs, regardless of their RMSD, display native-like binding energies. Our design process is fast and can rapidly cover large protein surfaces. In fact, used against the entire protein surfaces, it is capable to highlight interaction sites, similar to the previously published method MaSIF^20^. Besides capturing motifs of existing native complexes, our method can produce new binding motifs for a given site together with an assigned probability. We discuss in detail a real-world design scenario by targeting influenza’s hemagglutinin (HA) and the receptor binding domain (RBD) of SARS CoV-2. We believe that the presented new methodology will greatly contribute to development of new protein-protein interactions.

## Results and Discussion

### Workflow of the iNNterfaceDesign pipeline

iNNterfaceDesign is a framework consisting of several NN based models combined into one pipeline (Fig. 1a). First step of the framework is to design 6-residue backbone in the presence of target molecules using PepBB (peptide backbone). PepBB comprises three distinct stages each of which is carried out by separate attention-based neural network models (Fig. 1b). PepBB utilizes descriptors of the binding site only initially; during a flow of the data through the framework, the system accumulates predicted information about features of the motifs and sends them forward to the subsequent models.

**Figure 1.**
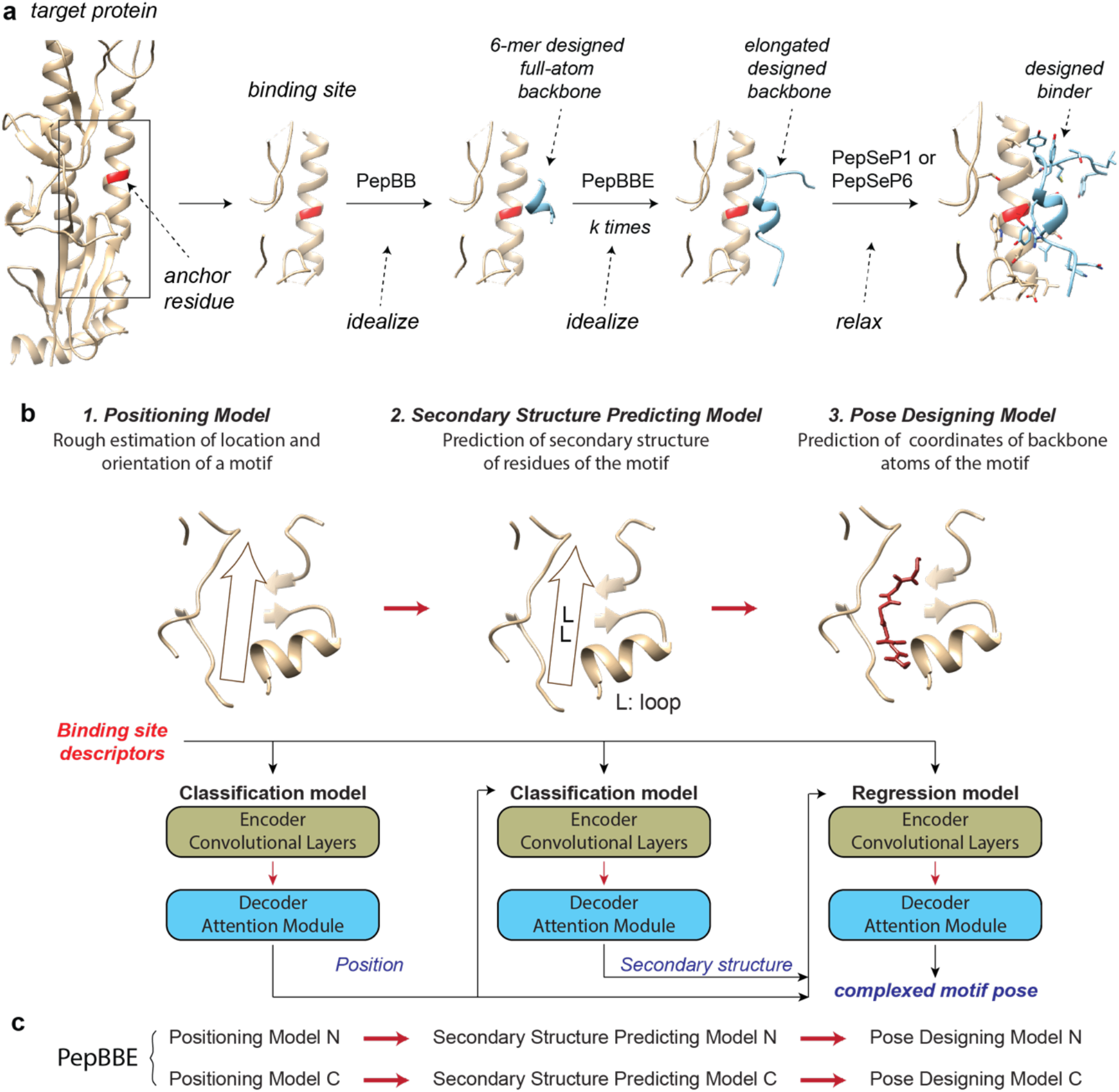
Overview of iNNterfaceDesign method. **(a)** General scheme of the method. **(b)** General scheme of PepBB and underlying deep learning models. **(c)** Scheme of PepBBE method elongating the designed peptides form C-and N-termini.

The targeted site on a protein is provided through a user-defined anchor residue (AR, Fig. 1a) or, if a larger surface area is targeted a list of anchor residues. A list of specific interface residues (IR_list) can be provided to focus within the AR defined area. A “rapid scan” version of our method of any surface array can be also applied to identify ARs; we demonstrate below that is procedure can identify preferential binding sites that overlap with natural protein-protein interactions sites. The first model of PepBB is a positioning model designed to generate approximate location and orientation of the backbone. The second model, called the secondary structure predicting model, produces as its name indicates a secondary structure sequence for two central residues based on the binding site descriptors and the output of the positioning model. The final step of the framework is prediction the concrete complexed 6-residue backbone fragment having all non-hydrogen backbone atoms represented by an array of distance maps defining XYZ-coordinates. Backbone atoms are then idealized using Rosetta. An additional option to swap the generated backbones with native backbones having close geometry from our *in-silico* library of 6-residue backbone fragments is available too. However, the backbones generated *de novo* for this study, except for Fig. 5.

Generated backbones can be extended using PepBBE (peptide backbone elongating) making them readily accessible for molecular grafting methods. PepBBE follows the same tasks as PepBB (Fig. 1c). For extension, the first three residues need to be able to replicate the position of last three residues of the initial designed backbone provided as input (Fig. 2b). The procedure of increasing of the length of the fragment can be repeated a few times on either end.

**Figure 2.**
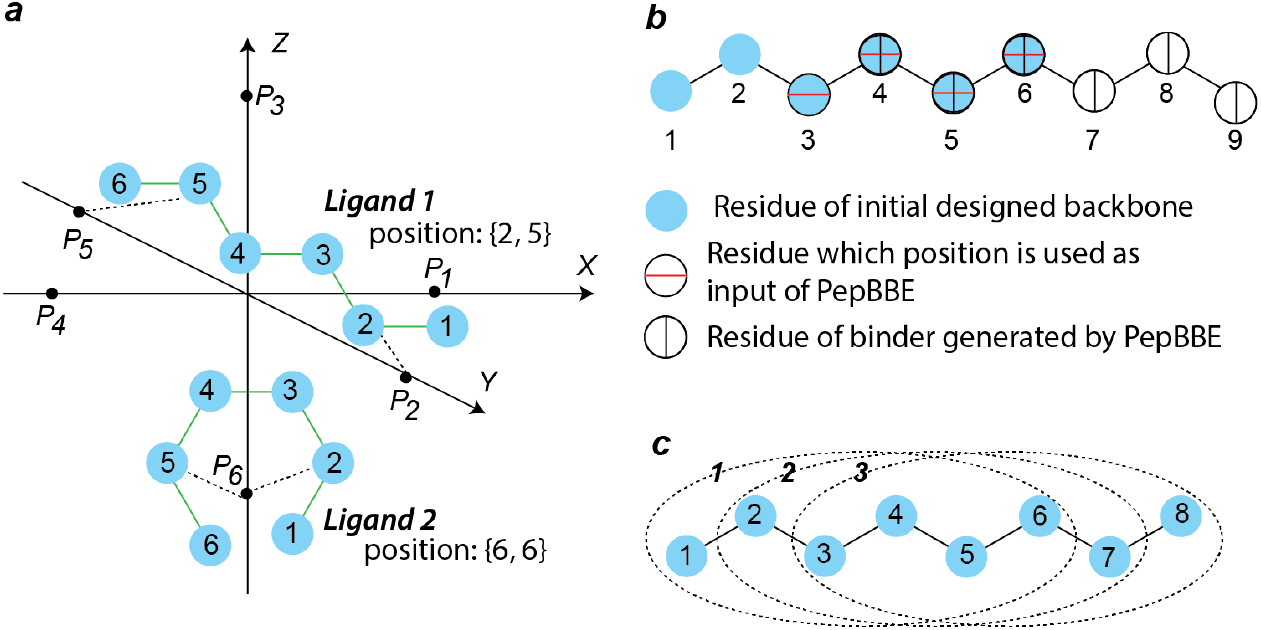
Illustration of key principles of PepBB and BepBBE methods. **(a)** Determination of a position of a peptide fragment: cartesian coordinate system with indicated points P_1-6_ on the x, y and z-axes, located at distance of 10 Å from the origin, which are used for defining the closest rays to second and fifth peptide residues. **(b)** Elongation of a designed backbone from C-terminus. **(c)** Three subsequent 6-residue fragments of native protein ligands which features are used as equally correct desirable outputs of the neural network models.

Amino acid sequence (AAS) design for the new backbones is then performed by the deep NN-based PepSeP1 or PepSeP6 methods^21,22^. These two methods were specifically developed for iNNterfaceDesign. The advantage of these deep neural network-based methods is that they do not need ideal positioning and final geometry of the backbone for the sequence design of the fragments; in fact, they can guide the refinement process. Biophysically correct geometries and bound conformation only have to be fulfilled in the final refinement step. We believe this is a major advantage over purely energy function-based methods which may reject a solution before being able to refine it.

### Performance of each deep neural network model in PepBB relative to native PPI

We evaluated performances of all models of PepBB on an independent test set. Test set consist of 24-48 residue binding sites complexed with 8-residue peptide ligand fragments, these interacting patches are cut out from native PPIs (Fig. S1). The samples, 1264 in total, are selected according to following main criteria: two out of central 6 residues of a native peptide ligand fragment (this fragment comprising 6 native central residues is referred as a core peptide fragment further) contribute to the binding with ΔΔ*G*_*i*_ > 0.5 REU and at least three of out of four central residues locate within 6 Å from a binding site when measuring distance between closest heavy atoms (side chains are considered), this distance threshold is denoted as *Thr*. Test set is divided into T*-*ho and T*-*he subsets, based on whether they are derived from homo- and hetero-oligomeric protein-protein interfaces correspondingly (more under S2.1-2.2.).

### Challenges to recapitulating native protein-protein interactions

One of the challenges to evaluate the performance and its ability to recapture native PPIs is determination of a starting point for the design of a new peptide ligand. The anchor residue can be the closest surface residue for more than one ligand residues simultaneously (Fig. S2). Therefore, there is a certain probability, that the predicted fragment would correspond to the experimental structure with shift of ± k residue(s), where k ≤ 6. To overcome this problem in part, we utilized 8-residue long native ligand fragments to compare our backbone fragments to; these 8-residue peptides can be represented as three 6-residue peptides (with a stride of 1 residue difference, Fig. 2c) all of which we use to evaluate performance of the method. Performances of separate models of PepBB method is presented in Table 1.

**Table 1.** Performance of models of PepBB method on test set (T) and its subsets. Performance of secondary structure predicting model (SS model) is measured by providing native positions and performance of pose designing model is measured by providing native positions and secondary structures of core peptide fragments here.

| Subset | Secondary structure | Positioning model |  |  |  |  |  | SS model |  | Pose designing model |  |  |  |  |
| --- | --- | --- | --- | --- | --- | --- | --- | --- | --- | --- | --- | --- | --- | --- |
| | | Number of Samples | RO3, % | RO3-1, % | RO3Rev, % | RO1, % | $N^3_{pos}$ 90% | RS1, % | RS2, % | RMSD4, Å | RMSD6, Å | RMSD6r, Å | RMSDh, Å | RevP, % |
| T-ho | α-helix | 338 | 65.68 | 25.44 | 1.18 | 47.93 | 2 | 76.92 | 78.55 | 3.91 | 4.66 | 4.24 | 2.49 | 27.81 |
| | β-sheet | 142 | 52.11 | 30.99 | 9.15 | 31.69 | 3 | 58.45 | 63.03 | 3.9 | 5.28 | 4.71 | 3.2 | 12.68 |
|  | loop | 208 | 51.44 | 32.69 | 7.21 | 28.37 | 3 | 57.21 | 59.86 | 4.04 | 5.39 | 5.12 | 3.27 | 14.9 |
|  | mixed | 185 | 44.86 | 41.62 | 7.03 | 31.35 | 4 | 49.19 | 66.22 | 3.67 | 4.94 | 4.75 | 3.1 | 9.73 |
|  | Overall | 873 | 55.67 | 31.5 | 5.15 | 37.11 | 3 | 63.34 | 68.96 | 3.89 | 5 | 4.63 | 2.92 | 18.44 |
| T-he | α-helix | 120 | 59.17 | 30.83 | 0.83 | 45 | 3 | 65.83 | 66.67 | 4.05 | 4.85 | 4.36 | 2.77 | 27.5 |
| | β-sheet | 77 | 46.75 | 35.06 | 5.19 | 35.06 | 4 | 53.25 | 55.84 | 3.34 | 4.44 | 4.34 | 3.21 | 5.19 |
|  | loop | 95 | 46.32 | 37.89 | 3.16 | 29.47 | 3 | 48.42 | 55.26 | 4 | 5.32 | 4.96 | 3.32 | 15.79 |
|  | mixed | 99 | 66.67 | 29.29 | 5.05 | 40.4 | 3 | 48.48 | 66.16 | 3.79 | 5.03 | 4.76 | 2.9 | 18.18 |
|  | Overall | 391 | 55.5 | 32.99 | 3.32 | 38.11 | 3 | 54.73 | 61.64 | 3.83 | 4.93 | 4.6 | 3.03 | 17.9 |
| T | α-helix | 458 | 63.97 | 26.86 | 1.09 | 47.16 | 2 | 74.02 | 75.44 | 3.95 | 4.71 | 4.27 | 2.56 | 27.73 |
| | β-sheet | 219 | 50.23 | 32.42 | 7.76 | 32.88 | 3 | 56.62 | 60.5 | 3.7 | 4.99 | 4.58 | 3.2 | 10.05 |
|  | loop | 303 | 49.83 | 34.32 | 5.94 | 28.71 | 3 | 54.46 | 58.42 | 4.03 | 5.37 | 5.07 | 3.29 | 15.18 |
|  | mixed | 284 | 52.46 | 37.32 | 6.34 | 34.51 | 3 | 48.94 | 66.2 | 3.71 | 4.97 | 4.75 | 3.02 | 12.68 |
|  | Overall | 1264 | 55.62 | 31.96 | 4.59 | 37.42 | 3 | 60.68 | 66.69 | 3.87 | 4.97 | 4.62 | 2.96 | 18.28 |

After the AR(s) have been defined, the positioning model starts by with predicting probable orientation and position of the newly designed peptide fragment as defined by the order number of rays (x, y, z, -x, - y, -z) of a coordinate system. The coordinate system originates at the AR, closest to Cα atoms of 2^nd^ and 5^th^ residues of the peptide fragment (Fig. 2a). Hence, there are six degrees of freedom according to number of the rays related to these two residue positions, resulting in 36 possible combinations from which the positioning model is guessing. Therefore, it converts this regression problem (since the actual number of locations and orientations is infinite) into a classification problem improving its predictive performance. However, peptide fragments having identical neighboring rays for both ends (Fig. 2a, ligand 2) can result in ‘reversed’ designs. This would mean the correct location but with the opposite orientation. This is mostly only relevant in context of recapitulating native interactions.

#### Performance of the positioning model

The position of the 6-residue peptide fragment output is defined by 2 numbers within an internal coordinate system. They are the numbers of the 6 points P ∈ (P1, P2,.. P6) (Fig. 2a) closest to the 2nd and 5th residues of the fragment. The prediction of the positioning model is labeled as true if it is matching with features of one of the adjacent protein ligand fragments (Fig. 2c). A few metrics are employed to evaluate the performance (Table 1):

a. RO3 is a ratio of true guesses to the total number of input samples.
b. RO3-1 is a ratio of guesses where only one number is predicted correctly, e.g., {4, 6} instead of target values {4, 2}, to the total number of input samples.
c. RO3Rev is a ratio of true guesses with reversed order of indices, e.g., {2, 4} instead of target values {4, 2}, to the total number of input samples; samples counted in RO3 are not considered. The resulted position corresponds to the reverse design.
d. RO1 is a ratio of true guesses for the core fragment (Fig. 2c, peptide 2) to the total number of input samples.
e. 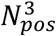 90% is number of guessed positions needed to have the native position among them given a 90% confidence interval. Positions were ranked according to their model probabilities.

As expected, the highest success rate is observed for helical fragments, which has an RO3 of about 59-65%. A high accuracy of 66.67% is achieved on fragments having mixed secondary structure on subset T-he, as well. RO3 measured on peptides with other types of secondary structures is about 44-55%, but higher rates of false results are partially compensated by substantially increased RO3-1 and RO3Rev. RO1 is lower than RO3 by about 15-20% and has similar dependencies on the secondary structure of the peptides. Our method allows to iterate through a few positions resulting in different poses of the peptide fragments. Therefore, we calculated the statistic metric 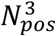 90% to estimate the optimal number of positions to try to get a native positioning among the predictions with a 90% of confidence level. The results indicate that two positions are enough for consideration in case of helical structures, but the number is 3-4 usually.

#### Performance of the secondary structure predicting model

The secondary structure predicting model predicts as its name indicated the secondary structure types of two central residues of the 6-residue peptide fragment. Two metrics are introduced to evaluate the performance of the secondary structure predicting model:

a. RS1 is a ratio of the peptides where identities for both residues are predicted correctly.
b. RS2 is a ratio of true guesses to the total number of the input residues.

Correct predictions are those which are matching with features of one of adjacent protein ligand fragments having the same position as provided to the model, the provided positions are positions of the core native fragment (Fig. 2c).

Secondary structures are predicted correctly for more than half of the peptide fragments, according to RS1 metric. The highest rates of the recovery are observed for helical fragments: RS1 is about 65-77%. Peptide fragments containing residues with different secondary structure types (mixed) are the hardest to predict: RS1 in around 49% in all test subsets. However, RS2 is substantially higher with 66% indicating that one of the points is indeed predicted correctly. For all other structure types (H/E/L), RS2 is similar to RS1. A confusion matrix was made up on base of RS2 metric (Figure S3). It visualizes the distribution of native and predicted secondary structure types of the residues. The most challenging sections are prediction of loop residues instead of regular structures; in case of native β-sheet fragments, the share of predicted loop residues is almost on par with the correct guesses.

#### Performance of pose designing model

As reported for the positioning model, we used 3 predicted 6-mer fragments to compare to a native 8-mer fragment (due to the challenge discussed above). We then measured the average distance between Cα backbone atoms of superimposed native and designed peptide fragments. The lowest RMSD among the 3 predicted fragments was reported (Table 1). All adjacent native protein ligand fragments of the training set were used as equally correct outcomes during training of the pose designing model by means of our custom loss function. Positions and secondary structures of native motifs are provided to the model to evaluate its performance here. We utilized several RMSD metrices for our performance evaluations:

a. RMSD6 is the RMSD between poses of designed and native closest ligands.
b. RMSD4 is the RMSD considering nonterminal residues only.
c. RMSD6r is the RMSD6 where RMSD for reversed designs (the fragments are on same location but N- and C-termini are flipped) was calculated on base of reversed numbering of the designed residues.
d. RMSDh is the RMSD between hotspot residues^23,24^ of native fragment and closest to the Cα-atoms in the designed peptides; they are defined as residues interacting with ΔΔG_i_ > 1 REU, unless otherwise specified.
e. RevP is the share of designed reversed poses.

The RMSD6 and RMSD4 values are 4.97 Å and 3.87 Å, respectively (Table 1). As expected, the results are sensitive the distance threshold (*Thr*) (Fig. S3). The shorter the distance between interfaces in the native complex, the lower is RMSD metrics, so RMSD4 is equal 3.3 Å at *Thr* of 3 Å. The difference between the 4 or 6-residue long fragment alignment is 1.1 Å and indicates that residues at the end of the peptides are harder to predict, probably due to absence of distance restrictions imposed on them. The performance of the model is similar for both homo- and hetero-oligomer subsets. RMSD6r is less than RMSD6 by 0.3-0.4 Å approximately. It is remarkable that RMSDh is considerably lower than both RMSD6 and RMSD6r, designed residues are within 2.5 Å of hot spot residues for 45% of our predictions (Fig. 3a).

**Figure 3.**
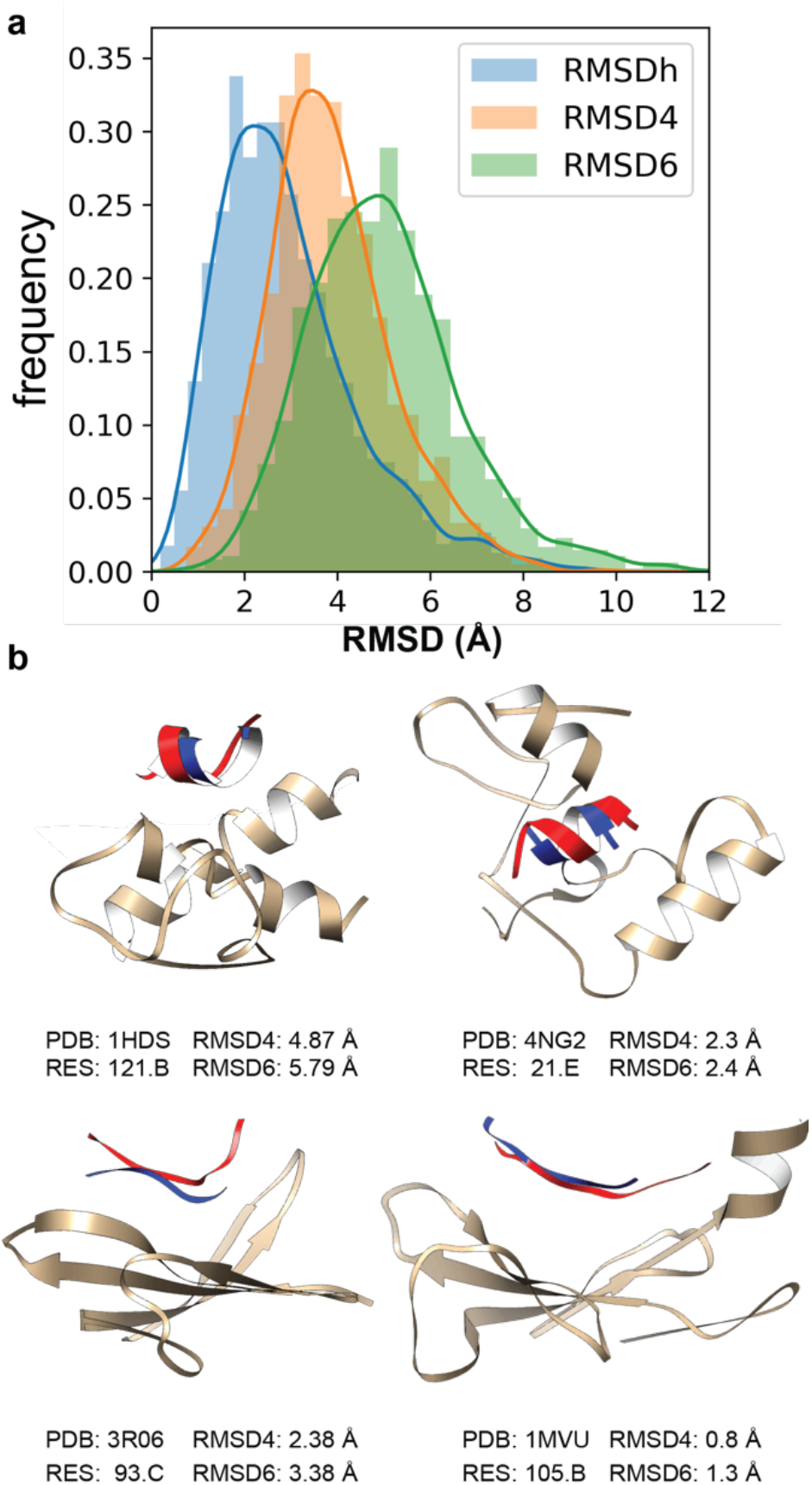
(a) Distribution of the RMSD values of the peptide fragments or hotspots with respect to native conformations. (b) Visualization of randomly selected complexes to illustrate RMSDs from test set with designed 6-mer backbones and native fragment 8-mer (red). RES is second residue of the native protein fragment in the original crystal structure.

Secondary structure of native fragments is the factor providing the highest impact to the results. There is a certain dependence between RevP and RMSD4 (Fig. S5): α-helices with the highest share of reversed poses (27.73%) show one of the highest RMSD values, whereas β-sheets have the lowest RevP and RMSD4. At the same time, α-helices have the lowest deviations when their poses are estimated by RMSD6r and RMSDh (Table 1). High values of RMSD4 and RMSD6 in case of α-helices are due to their more compact shape in which both ends of the fragments are close to the same coordinate axis (Fig. 2a), making it more challenging to determine their orientation.

Fig. 3b demonstrate the quality of the generated backbones and the ability of the method to generate relevant backbone conformations for them. It should be mentioned that 6 poses (0.5%) of test set were designed with one or more missing heavy atoms. We do note that the method generates occasionally distorted helix backbones (Fig. S6) which should be filtered for future design applications. The performance of the extension design model PepBBE is discussed in detailed under S1.2.

### Performance of all three models combined into PepBB

For each of the test binding sites, 12 designed backbones were generated by using four most probable positions and three most probable secondary structure sequences for each of the positions. The most probable output pose is termed as “Top pose” while the pose with the lowest RMSD4 relative the native fragments is termed as “Closest pose”. Results of comparison of the designed backbones with native fragments using metrics as for estimation of performance of pose designing model are reported in Table 2. RMSD4 and RMSD6 are equal to 5.52 and 6.78 Å on the top poses of test set, respectively. This lower accuracy on the top poses is due to frequent generation of reverse oriented designs which share is close to 40%. The percentage of reversed designs is lower on hetero-oligomeric PPIs. RMSD6 and RMSD4 on the closest poses of set T are equal to 5.0 Å and 3.77 Å, correspondingly. Analysis shows that values of RMSD4 and RMSD6 on top poses is about 30% worse than on the closest poses. We illustrate examples in Fig. 3b and Fig. S14 to highlight that an RMSD4 of around 5 Å and higher of a single 6-mer fragment can be very close aligned to the native interactions and can capture its hotspot interactions.

**Table 2.** Summary of performance of PepBB method on test set and comparison of its results with performance of Docking Protocol of Rosetta upon docking of native peptide ligands in the latter case.

| Secondary structure | PepBB |  |  |  |  |  |  |  |  |  | Docking Protocol |  |
| --- | --- | --- | --- | --- | --- | --- | --- | --- | --- | --- | --- | --- |
|  | Top poses |  |  |  |  | Closest poses |  |  |  |  | Top poses | Best poses |
|  | RMSD4, Å | RMSD6, Å | RMSD6r, Å | RMSDh, Å | RevP, % | RMSD4, Å | RMSD6, Å | RMSD6r, Å | RMSDh, Å | RevP, % | RMSD6, Å |  |
| α-helix | 5.34 | 6.23 | 5.62 | 3.48 | 40.39 | 3.79 | 4.83 | 4.62 | 2.77 | 19 | 5.89 | 3.05 |
| β-sheet | 5.7 | 7.38 | 5.97 | 4.13 | 40.18 | 3.71 | 5.07 | 4.81 | 3.26 | 10 | 5.87 | 2.80 |
| loop | 5.9 | 7.28 | 6.43 | 4.23 | 45.21 | 3.94 | 5.26 | 5.12 | 3.35 | 9 | 5.77 | 3.00 |
| mixed | 5.3 | 6.67 | 5.95 | 3.75 | 34.51 | 3.61 | 4.95 | 4.81 | 3 | 10 | 5.73 | 2.96 |
| Overall | 5.52 | 6.78 | 5.95 | 3.84 | 40.19 | 3.77 | 5 | 4.82 | 3.05 | 13 | 5.82 | 2.97 |

The closest poses are sensitive to the applied *Thr* in the same way as the pose designing model is on its own: the performance decreases with an increase of the distance between interfaces of the native complexes, especially in the interval of 3-4 Å (Fig. S9). However, a decrease of RMSD values is less pronounced for the top poses (Fig. S9). Overall, RMSD for non-terminal residues of the top poses is around of 3.0-5.5 Å; the RMSD for all 6 residues is 5.0-7.0 Å. We cannot fairly compare these results with performance of docking protocols directly due to difference in length of peptides and origin of samples in benchmark sets. However, commonly used docking algorithms ^25–27^ report RMSDs exceeding 10 Å for at length of peptides 9-12 residues. PepBB is able to approximate the docked conformation of the native protein ligand fragments almost as well as a docking protocol. This is remarkable, as PepBB has to predict the complete backbone and docking conformation whereas the backbone for the docking protocols of the native ligands is not perturbed. To evaluate this in more detail, we docked native peptide fragments to the binding sites using the Rosetta Docking Protocol^28^. We generated 500 docked poses among which the highest scored (=top) poses and the best poses (=closest) to the native structures were selected. The results are reported in Table 2. The accuracy of docking results is only on average 1.5 Å better despite the advantage of the knowledge of the native peptide sequence and backbone conformation.

The sequence was designed by PepSeP6, a deep learning model producing 6 different AAS per fragment. Average binding free energies Δ*G*_*bind*_ of the top and the closest poses of the peptide fragments, after generating of AAS and selecting most energetically favorable designs, are -19.5 REU and -19.8 REU, respectively (Fig. S10). However, the difference between the energies of the top and closest poses can be substantial in case of a generated backbones is twisted geometry; this occurs occasionally (Fig. S6). Average binding affinity of the native fragments of the test set, relaxed with the same constraints as the designed peptides, is -19.7 REU. Hence, the difference between the energies of the native and the closest designed peptide ligands is very close. Comparison of amino acid distributions in the native and the designed peptide ligands of test set is presented in Fig. S11. Results presented in Table S4 indicates that 51-59% of the native interacting binding site residues and 58-67% of the hotspot residues bound with the designed motifs in a comparable way, structurally or energetically, depending on poses and the test subsets.

### Case study: Targeting influenza’s hemagglutinin and the receptor binding domain of SARS-CoV-2

To demonstrate two example applications, we targeted influenza’s hemagglutinin (HA) and the receptor binding domain (RBD) of the SARS CoV-2 virus. All crystal structures for HA or RBD were taken out of the training set to mimic an application to a previously “unseen” crystal structure. As the generation of motifs and design is rather fast, we performed a “rapid scan” to quickly generate motifs covering the entire surface of the target molecule. For the rapid scan, motifs are generated with all surface residues serving as an anchor residue (Fig. 4, 5). We designed six top poses for each anchor residue using PepBB combined with PepSeP1. The average binding affinities 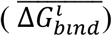 between the six motifs of the anchor residue *i* and the receptor are calculated and assigned to the anchor residues. A threshold of -10 REU is applied for 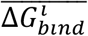, surface anchor residues which peptide ligands display affinity lower than the threshold (Fig. S12b) can be selected for further design. Concrete examples of further application of iNNterfaceDesign on HA and comparison of the designs with native protein ligand fragments from antibodies are discussed in SI 1.4. To generate binding proteins, the motifs can be grafted into our previously developed scaffold database^29^ and designed as small protein fold (Fig. S15) or potentially grafted into an antibody fragment framework as a CDR3. Alternatively, structures with matching secondary structures that could sustain the grafting of a designed motifs could be used.

**Figure 4.**
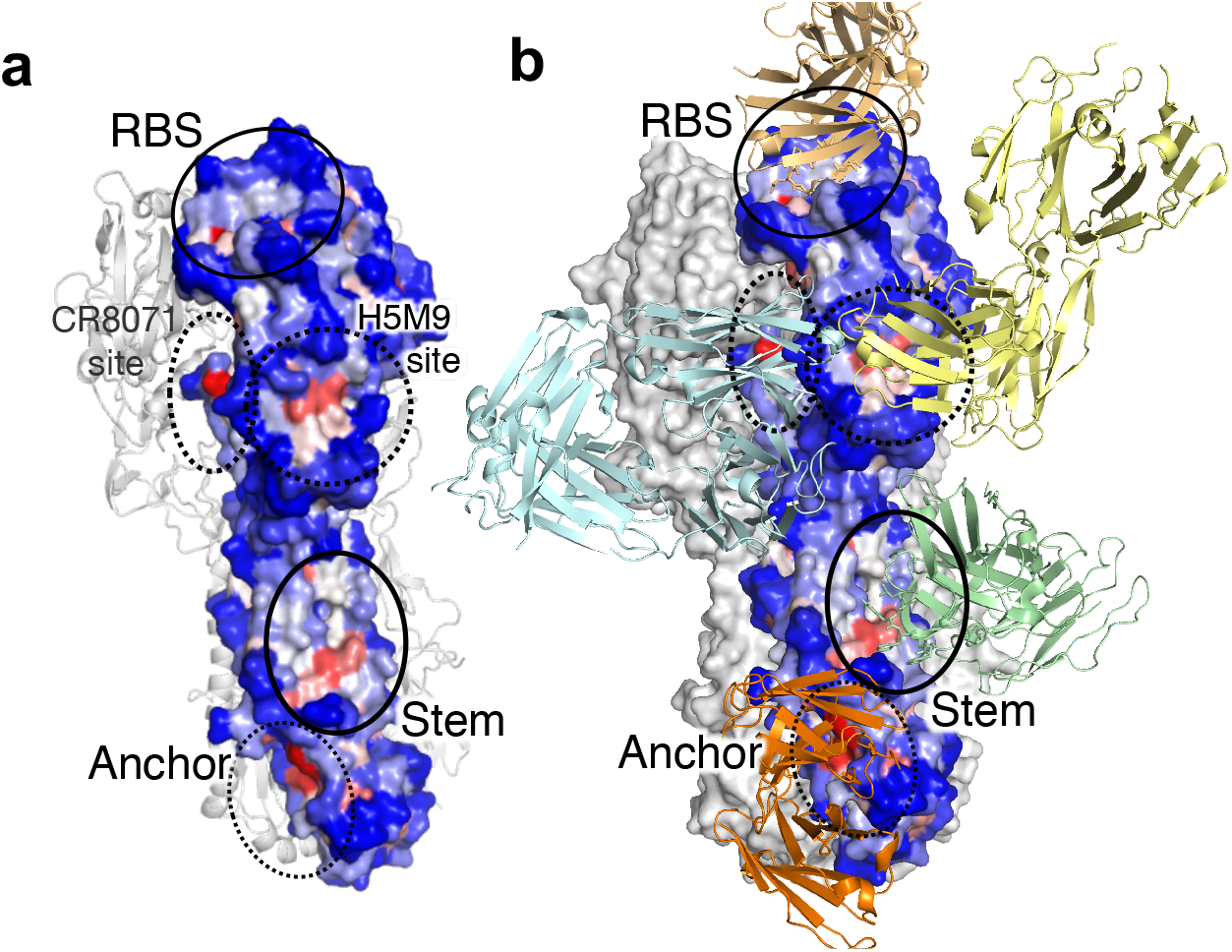
Binding sites on surface of H3 Influenza hemagglutinin (code: 3ZTJ) according to results of iNNterfaceDesign. (a) Privileged binding sites identified via our rapid motif scan which designed motifs against the entire surface of the hemagglutinin monomer; residues are colored according to 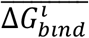 with red being low binding energies, and blue none to rather weak (high) energies. (b) Stalk binding antibody FI6 (3ztj, light green), anchor antibody (7t3d, orange), receptor binding site S139/1 or C05 (4gms, 5umn). We also included a superposition of H5M9 bound to H5 (A/goose/Guangdog/1/96), CR8071 bound to Influenza B HA (palecyan, 4fqj) as they illustrate neutralizing binding sites on broadly reactive antibodies to a site our algorithm highlighted.

**Figure 5.**
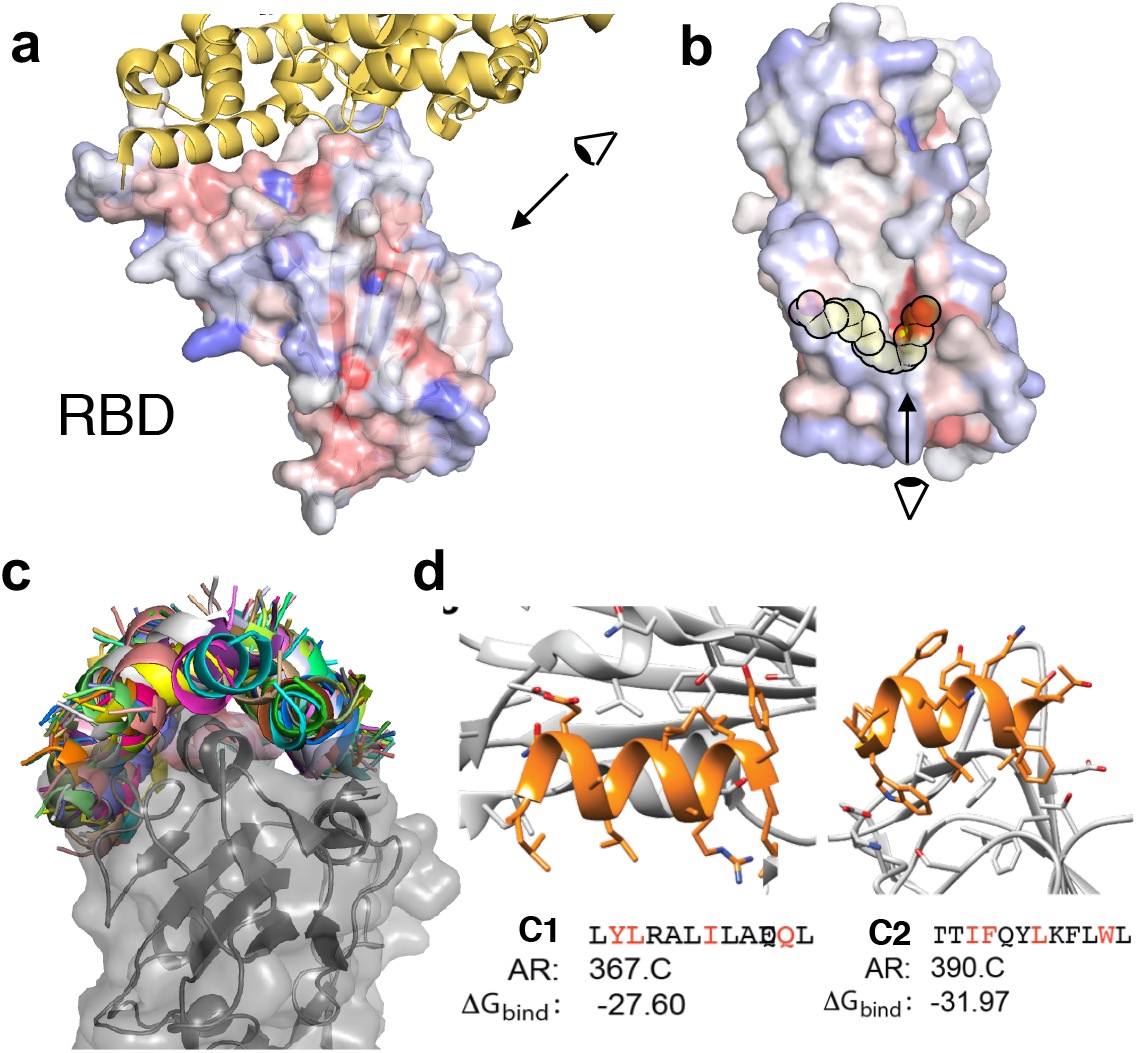
Performance of iNNterfaceDesign in design of motifs for SARS-CoV-2 (code: 6W41). (a) Anchor residues (AR) determined for SARS-CoV-2 and colored according to 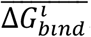. Co-crystal structure with the recptor Ace2 (yellow, 6m0j). Eye indicates perspective for Fig. 5b. (b) Linoleic binding site provides a potential excellent epitope for binding to the backside of the RBD. Eye indicates perspective for figure 5c. (c) Overview of generated 12-mer helical motifs targeting parts of the RBD domain (d) Visualization of 2 random motifs (c1, c2) interacting with the target with Δ*G*_*bind*_ ≤ -25 REU. Red letters in AAS corresponds to residues with contributions ΔΔ*G*_*i*_ ≥ 2.5 REU to binding.

When coloring the target structure based on 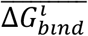 per anchor residue, we can identify preferred binding sites, similarly as deep learning method MaSIF^20^ which uses predefined geometric feature for detecting potential epitopes. However, iNNterfaceDesign method was not trained to identify preferential binding sites, it was trained on complexes and thereby learned what defines a privileged binding site without explicit implementation of general descriptors. Using our rapid scan mode, we see preferential interaction sites with the receptor binding sites for HA, the conserved stem region as well as some areas within the trimeric interface (Fig.4). Interestingly, we can also see another interaction site to which the H5M9 antibody binds (binding to most influenza B strains)^30^. While there is currently no antibody reported (at least a co-crystal structure) targeting H3 of influenza A, this could be an interesting target site. Furthermore, we see excellent recapitulation of antibody interactions of the broadly neutralizing antibodies (bnAb) 5j8 and FI6v3 with HA (Fig. S13, 14) demonstrating that the algorithm is indeed general and can next to longterm naturally evolved protein-protein interactions also reproduce the interactions of antibody loops with their immunogen.

For the RBD domain, we can see binding hotspots for the receptor binding site with Ace2 (Fig.5a) but also inside the linoleic acid binding site (Fig. 5b). We selected 53 anchor residues after an initial scan to design 6-Residue helical fragments (Fig. 5c). The helices were elongated using pepBBE and AAS were designed with PepSeP6. The iNNterfaceDesign pipeline produced numerous of designs (more than thousands in total per epitope). The number considerably decreases after application of different filters checking designs, but even after that, the method provide enough variety of backbones for one sided design (Fig. 5b, 2714 in total), especially if PepSeP6 is applied a few times for more diverse pool of AAS (method details are in ^21^). Depictured motifs (Fig. 5d) demonstrate that the method succeeds in generating of backbones and judicious placement of residues, including voluminous ones. Based on residues with ΔΔ*G*_*i*_ ≥ 2.5 REU, all motifs had hotspots incorporated (full results of alanine scanning ^31^ are summarized in Table S5).

## Conclusions

We have developed a fundamentally new approach to design protein-protein interface motifs. It is inspired by computer vision and uses several deep learning models to design new protein interaction fragments from scratch. Our design workflow consists of 3 methods within the iNNterfaceDesign pipeline. The first method, PepBB, predicts initial 6-residue backbone fragments of peptide ligands; second PepBBE elongates the initial backbones and then the third model, PepSep1 or PepSep6^22^ produces amino acid sequences for the designed motifs. We discussed the performance of the core part PepBB, which is the most crucial starting component in detail. Itself consists of 3 NN-based models with the first one predicting the rough location of the new peptide fragment given an anchor residue; second, it predicts its secondary structure and then the third model produces the atomic details of the new fragment. Fragments are then designed with our PepSeP1 algorithm which decides on the amino acid identities – also performed by a deep learning model. We demonstrate its performance by comparing it to a test set of homo- and heteromeric interactions of 179 proteins which have a diverse combination of secondary structures within their interfaces. Our new methods are capable of reproducing native-like interactions from our test set, including prediction of identities of hotspot residues and their positioning with high accuracy. We present two real-world scenarios in which we targeted H3 hemagglutinin and the RBD of CoV-2. For HA, we demonstrate that our algorithm is capable to reproduce antibody-antigen interaction loops, reproducing both backbone conformation as well as hotspot residues. Our “rapid scan” version of the methods allows to highlight favorable binding epitopes and facilitates thereby also the identification of a starting point (anchor residues). In case of HA, we can see the epitopes to which broadly neutralizing antibodies bind, and for RBD, we see how the receptor binding site is an “attractive” epitope as well as the linoic acid binding site. Therefore, our method also provides a new tool for the identification of PPI sites, similarly to the previously reported method MaSIF^20^. We believe our algorithm is generally applicable for the challenging problem of designing PPIs from scratch and will be a highly useful tool for the field of protein engineering and design, including the antibody engineering field.

## Methods

### Input and output data of the neural networks

There are seven types of input arrays provided to the neural networks:

1. Input 1 of size 48 × 6 × 2 consists of two distance maps of size 48 × 6 containing distances between backbone atoms of 48 binding site residues and 6 reference points which are similar to points P_1-6_ (Figure 2a) but located at a distance of 6 Å from the origin; Cα and O backbone atoms are used for the distance maps.
2. Input 2 is an array of size 48 which contains secondary structure sequence of the residues of the binding site.
3. Input 3 is an array of size 48 which contains AAS of the residues of the binding site.
4. Input 4 is an array of size 48 which distinguishes core, surface and interface residues (if IR_list is provided) of the binding site by different values (0.2, 0.6 and 1, respectively).
5. Input 5 is an array of size 2 which contains position of the peptide ligand predicted by the positioning model.
6. Input 6 is an array of size 2 which contains secondary structure sequence for two central residues of the peptide ligand predicted by the secondary structure predicting model.
7. Input 7 of size 4 × 6 × 2 consisting of two distance maps of size 4 × 6 containing distances between Cα and O backbone atoms of residues 1-4 and 5-8) of native 8-mer protein ligand fragment (Fig. 2c) till 6 reference points mentioned in description of input 1. They are used for PepBBE models elongating C- and N-termini correspondingly.

Zero padding is used if the number of the residues of the binding site is less than 48 in case of inputs 1-4.

Interface residues were used instead of anchor residue sometimes in this study. These are the pocket residues which are the closest to the native peptide fragment when the distances are measured taking into account all heavy atoms of the pocket; the lists, IR_lists, consist of the closest 8-25 residues (the total number is random) with random 2-5 residues deleted from the lists. The elements of random selection are due to frequent absence of information about exact interface residues in real application, one can guess them approximately only. Two-thirds of input 4 arrays of the training set do not point out interface residues and one-third of the inputs do.

First four input data are provided to the positioning model of PepBB, an output of which, output 1, is a vector of 36 probability values. Input 5 is quotient and remainder when index of the element with the maximum probability from output 1 is divided by 6. The secondary structure predicting model, based on 5 inputs, generates output 2 representing probabilities for secondary structure sequence of two central residues of the backbone.

The pose designing model is fed by all six input and produces output 3-4. Output 3 contains a predicted complexed backbone of the motif in a form of Cα atoms trace. Output 4 is an array of size 6 × 6 × 4 consisting of four distance maps corresponding to all heavy backbone atoms of the peptides. These maps are used for creation of full-atom model of the backbone conformation.

Inputs and outputs of PepBBE are like those of PepBB, but all its models are fed by input 7 additionally.

### Architecture of Developed Neural Networks

An architecture of all models is very similar and loosely based on an TensorFlow tutorial ^32^ about how to implement models generating image captions with visual attention ^9^, specifically the decoder part. The encoders are based on convolutional layers. The neural network was built in TensorFlow using Keras API. More details under SI 2.4 -2.6.

### Recovery of Cartesian coordinates of backbone atoms from the outputs of the neural network

Recovery of backbone is based on two distance maps in output 3 defining distances between Cα atoms and points P_1-6_ as in (Fig. 2a) but located at distance 6 Å from the anchor residue. The atoms are predicted in direct and reversed order in two maps for increasing a probability of recovery of all residues, aligning of two resulted poses produce more accurate motifs also. Four shortest predicted distances till points P_1-6_ are selected for each atom. Trilateration procedure is carried out based on three closest reference points and the distances, the result is two points symmetric by the plane containing atoms used for the trilateration; then the point whose distance till the 4th reference point is similar to the predicted distance has been chosen. The full-atom backbone conformation is generated based on distances from output 4. This fragment is aligned with the Cα atoms trace recovered from output 3 earlier by means of rotation and translation operations; RMSD between Ca atoms is minimized using Kabsch algorithm.

### Assessment of experimental and predicted motifs

#### Refinement and calculation of binding energies of the designed complexes

The target binding sites were relaxed without peptide ligands. AAS predicted by PepSeP1 or PepSeP6 methods were mapped to designed idealized backbones and attached to the binding sites. Optimization of side chain conformations of all residues of the complex and subsequent refinement of poses of the peptide ligands was done using three times using FastRelax protocol ^33,34^ over 300 step while applying harmonic constraints (SD=3.0 with width parameter of 2.0 Å) Binding free energies of the complexes were estimated using InterfaceAnalyze^35^ with repacking chains after separation. All stages of the refinement were carried with FastRelax. The structure with the lowest score out of three results was selected for the next operation. The Rosetta scoring function ref15 was used for all calculations.

#### Docking of native peptide ligands using Rosetta’s Docking Protocol

Side chains of target binding sites and native peptide ligands were relaxed separately by deleting residues of counterpart from initial complex. The structures were attached back without changes in positions of backbone atoms relative native complexes after. Then Docking protocol, generating 500 poses, was applied as follows:

<DockingProtocol name=“docking” docking_score_high=“sfxn_std” dock_min=“1” docking_local_refine=“0” ignore_default_docking_task=“0” />.

### Code availability

The source code of the method is available at https://github.com/strauchlab/iNNterfaceDesign

## Notes

### Competing Interest Statement

The authors have declared no competing interest.

